# The Effect of Phylogenetic Uncertainty and Imputation on EDGE Scores

**DOI:** 10.1101/375246

**Authors:** K. Bodie Weedop, Arne Ø. Mooers, Caroline M. Tucker, William D. Pearse

## Abstract

Faced with the challenge of saving as much diversity as possible given financial and time constraints, conservation biologists are increasingly prioritizing species on the basis of their overall contribution to evolutionary diversity. Metrics such as EDGE (Evolutionary Distinct and Globally Endangered) have been used to set such evolutionarily-based conservation priorities for a number of taxa, such as mammals, birds, corals, amphibians, and sharks. Each application of EDGE has required some form of correction to account for species whose position within the tree of life are unknown. Perhaps the most advanced of these corrections is phylogenetic imputation, but to date there has been no systematic assessment of both the sensitivity of EDGE scores to a phylogeny missing species, and the impact of using imputation to correct for species missing from the tree. Here we perform such an assessment, by simulating phylogenies, removing some species to make the phylogeny incomplete, imputating the position of those species, and measuring (1) how robust ED scores are for the species that are not removed and (2) how accurate the ED scores are for those removed and then imputed. We find that the EDGE ranking for species on a tree is remarkably robust to missing species from that tree, but that phylogenetic imputation for missing species, while unbiased, does not accurately reconstruct species’ evolutionary distinctiveness. On the basis of these results, we provide clear guidance for EDGE scoring in the face of phylogenetic uncertainty.

## Introduction

Evidence from the fossil record and present-day studies argue we are in the midst of, or entering, a sixth mass extinction (Barnosky et al., 2011; Ceballos et al., 2015), such that more populations than ever are declining and species face heightened danger of extinction (C. D. Thomas et al., 2004; Wake & Vredenburg, 2008). Habitat destruction (Brooks et al., 2002), invasive species (Molnar, Gamboa, Revenga, & Spalding, 2008), climate change (Pounds et al., 2006), and disease (Lips et al., 2006) are some of the leading causes of species declines globally. Conservation biologists seek to reduce these detrimental effects on species populations, but in reality they have limited resources with which to do so. This challenge, termed the “Noah’s Ark problem” (Weitzman, 1998), has driven conservation biologists to identify different ways by which to prioritize, or triage, their resource allocation (Bottrill et al., 2008).

Conservation triage, like all sound decision-making, requires an index with which to quantify the relative urgency or importance for conservation among a set of options. This allows scientists and policy-makers to use data to quantify need and inform conservation decision-making and management activities. One triage strategy uses the EDGE metric to identify and prioritize species that are Evolutionarily Distinct and Globally Endangered (Isaac, Turvey, Collen, Waterman, & Baillie, 2007). Evolutionary Distinctiveness (ED) measures the relative contributions made by each species within a particular clade to phylogenetic diversity, assigning each branch length equally to all the subtending species (Isaac et al., 2007; D. Redding, 2003). Global Endangerment (GE), assigns numerical values to each of the World Conservation Union (IUCN) Red List Categories. As species become increasingly threatened and are placed into categories of increasing concern (*e.g.* from Vulnerable to Endangered), the GE numerical value increases. A species’ EDGE score is an aggregate value intended to equally reflect a species’ evolutionary distinctiveness and conservation status (even if it does not always in practice; see Pearse et al., 2015).

Usage of the EDGE metric has expanded greatly. First used to prioritize global mammals (Isaac et al., 2007), EDGE scores are now available for a variety of taxonomic groups, including amphibians (Isaac, Redding, Meredith, & Safi, 2012), birds (Jetz et al., 2014), corals (Curnick et al., 2015), squamate reptiles (Tonini, Beard, Ferreira, Jetz, & Pyron, 2016), sharks (Stein et al., 2018), and all tetrapods (Gumbs, Gray, Wearn, & Owen, 2018). Related metrics are also now available, each subtly emphasizing different things, such as the expected contribution of each species to future phylogenetic diversity (HEDGE, I-HEDGE; Jensen, Mooers, Caccone, & Russello, 2016; Steel, Mimoto, & Mooers, 2007) and our uncertainty over a species’ future (EDAM; Pearse et al., 2015) The development and expansion of EDGE-like metrics mirrors progress in other areas of conservation biology, and the likelihood of success in conservation (McBride, Wilson, Bode, & Possingham, 2007; Wilson et al., 2007), the relative cost of certain interventions (Naidoo et al., 2006), and complementarity of interventions (Myers, Mittermeier, Mittermeier, Da Fonseca, & Kent, 2000; Pressey, Humphries, Margules, Vane-Wright, & Williams, 1993) can now be considered in its calculation. The EDGE index was developed explicitly with the intention of informing conservation triage, and is now the basis of the global EDGE of Existence Program (http://www.edgeofexistence.org/). The successful application of EDGE highlights the potential for phylogenetic conservation prioritization metrics to provide actionable insights while quantitatively measuring the evolutionary history a species represents. Nonetheless, almost every application of an EDGE-type approach must address uncertainty resulting from missing data.

Missing data can affect EDGE scores in several ways. First, the IUCN identifies some species as Data Deficient (IUCN, 2001, 2008), which affects the GE component of a species’ EDGE score. Fortunately, the IUCN provides guidance for using any available contextual data to assign some threat status to such species. A number of studies illustrate how to assign threat categories to Data Deficient species, which in turn should reduce the uncertainty in GE (Butchart & Bird, 2010; Dulvy et al., 2014; Good, Zjhra, & Kremen, 2006; Morais et al., 2013). The issue of missing phylogenetic data is arguably more complicated because not only does the focal species have no ED score, but its absence from the phylogeny may affect the ED scores of related species. Species of conservation concern are almost by definition rare, and frequently lack sufficient DNA (or even morphological) data to be placed with certainty on a phylogeny. In most cases, taxonomic information rather than sequence data alone has been used to place species in the tree of life when constructing EDGE lists (see Collen et al., 2011; Curnick et al., 2015; Forest et al., 2018; Gumbs et al., 2018; Isaac et al., 2012; Isaac et al., 2007; Jetz et al., 2014; Stein et al., 2018). Yet, to our knowledge, there has been no systematic study of the effect of imputation on species’ EDGE scores, despite this practice having received attention in other areas of comparative biology (Kuhn, Mooers, & Thomas, 2011; Rabosky, 2015; G. H. Thomas et al., 2013). Thus We do not know how accurate EDGE scores are when species are missing, or when species are added to phylogenies by imputation, nor do we know how accurate EDGE scores for imputed species might be. As interest in using EDGE-type measures and phylogenies for conservation triage grows, the need for consensus on how to resolve cases of phylogenetic uncertainty becomes increasingly urgent.

Here we attempt to quantify the effect of one sort of phylogenetic uncertainty - the effect of missing species on EDGE rankings - and assess the degree to which subsequent imputation affects the accuracy of EDGE scores. We do so by simulating phylogenies and then removing species either at random, or with bias, across those phylogenies. By contrasting the ED scores of the species before and after the loss of other species from the phylogenies, we measure the impact of missing species on ED scores. We then assess the extent to which phylogenetic imputation can accurately estimate the EDGE scores of missing species in simulated data. We also examine the extent to which such imputation affects the scores of species for which we have data. In doing so, we hope to provide clear guidance as to the applicability of phylogenetic imputation as a solution for species missing phylogenetic data. From our results, we argue that species’ ED values are remarkably robust to missing species, and that phylogenetic imputation does not reliably reconstruct the true ranking of those missing species.

## Methods

We use a simulation approach to test the effect of having missing species on a phylogeny (through species removal from simulated phylogenies) and then imputing species for species’ ED (Evolutionary Distinctiveness) scores. We focus exclusively on the ED-component of the EDGE metric, since uncertainty in species GE scores has already been addressed by the IUCN’s proposal to assign Data Deficient species scores (IUCN, 2001, 2008). Because EDGE is the product of both ED and GE components, even perfectly accurate GE values could be associated with imperfect EDGE scores if the ED scores were inaccurate.

All trees (both starting and imputed) were simulated under a pure-birth Yule model using ‘gieger::sim.bdtree’ (setting parameters b=1 and d=0; Pennell et al., 2014). This model was chosen because it is the simplest model possible: speciation rates are constant across the entire tree of life and there is no extinction. We acknowledge that more complex and/or biologically realistic models of diversification could potentially improve the performance of imputation. However, we suggest that imputation under a simple model that is identical to that used to simulate the data is a low, and fair, benchmark for a method to meet. We used ‘caper::ed.calc’ to calculate ED values (Orme et al., 2013). All simulations and analyses were performed using R (version 3.4.0; R Core Team, 2017), and we performed 100 replicate simulations of each parameter combination. All our analysis code is available online (https://github.com/bweedop/edgeSims).

### The impact of missing species on EDGE scores

Our first set of simulations assess the impact of missing species data on the ED scores of remaining species, considering data missing either in a random or phylogenetically-biased fashion. We simulated phylogenies of different sizes (number of species: 64, 128, 256, …, 2048, 4096) and then removed constant fractions of tips from the tree (0%, 1%, 2%, …, 19%, …, 99%). To simulate species missing at random throughout the phylogeny, we used ‘sample()’ to select the relevant fraction of species (rounded to the nearest whole number) without replacement. To remove species in a phylogenetically-biased manner, we used Felsenstein (2005)’s threshold model. We simulated a trait under a constant rate Brownian-motion model (σ=0.5, starting root value = 1) using geiger::sim.char’ (Pennell et al., 2014). Species were then removed from the tree if their simulated trait was in the upper quantile matching the fraction of species to be removed. For example, if 10% of species were to be removed from the tree, the species with the highest 10% of values would be removed. This results in closely related species being removed more often than expected by chance.

To quantify the effect of these manipulations, we calculated the ED values of species that are not removed from a tree both before and after removal. We then correlated these ED scores: if missing species do not affect ED values of the remaining species, we would expect a strong, positive correlation between the ED scores of the remaining species calculated before and after species were removed from the phylogeny. We emphasize that species removed from the phylogeny are omitted from this comparison. We outline our approach in figure 1.

**Figure 1:**
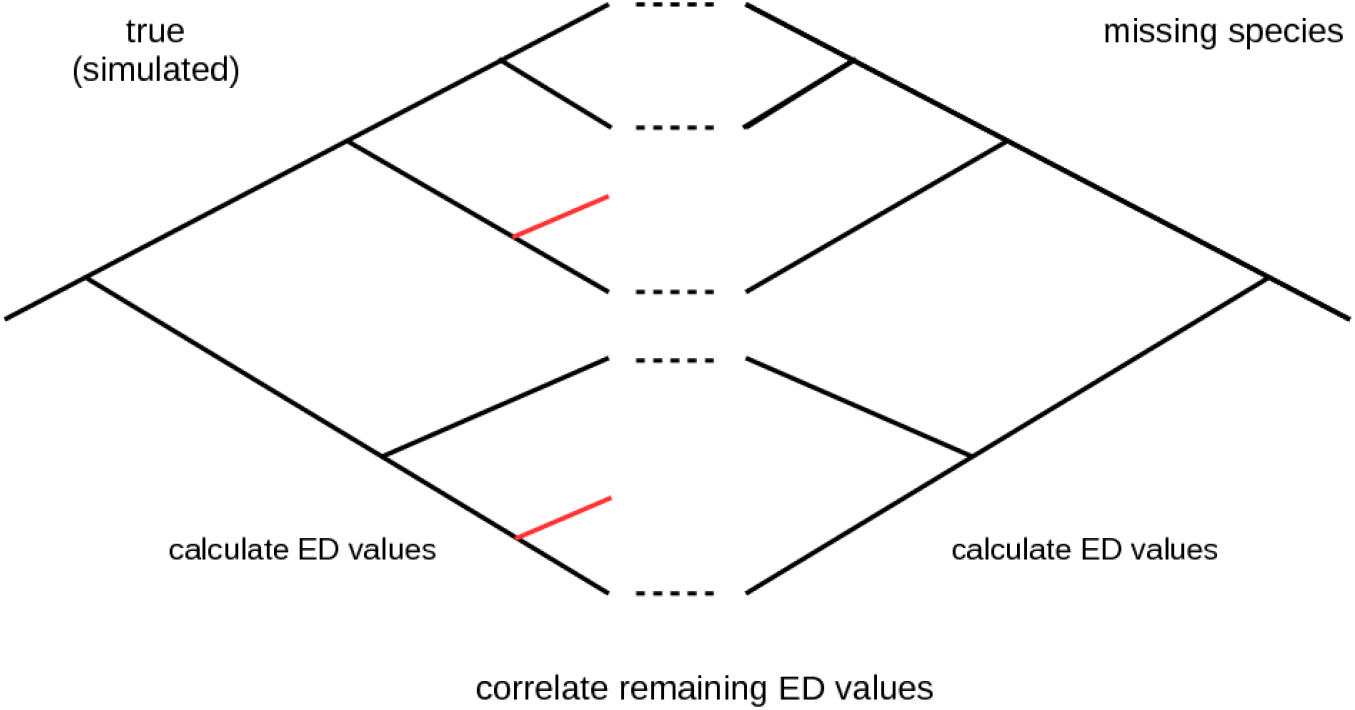
Conceptual overview of the missing-species simulations in this study. The simulated tree on the left is the true tree prior to removal of missing species. On the right is the same tree after missing species have been removed. Species that are removed are shown in red. To compare the ED values of the remaining species, we correlate their ED values before (left) and after (right) removal of the missing species. Dashed lines can be seen for the species which would have ED scores compared.

### The impact of phylogenetic imputation on EDGE scores

Our second set of simulations tested the impact of imputation on ED scores within an imputed clade. We used relatively small clades (5, 6, 7, …, 30, 31, 32 species) from phylogenies of different sizes [128 (2^7^), 147 (2^7.2^, 168 (2^7.4^), …, 776 (2^9.6^), 891 (2^9.8^), 1024 (2^10^) species). We first randomly selected a clade to be removed from the ‘true’ tree and then simulated a new phylogeny of the same size as the removed clade. This newly simulated clade was generated under the same pure-birth model as the original phylogeny. We then placed the newly simulated clade in the full phylogeny, in the same location as the removed clade. If a newly simulated clade was so old that it was not possible to graft it into place, we discarded that clade and simulated another. In an empirical study the model of evolution under which the phylogeny had evolved would have to be estimated, which is an additional source of error not considered here. We simulated each combination of clade and total phylogeny sizes 100 times. An overview of our approach is given in figure 2.

**Figure 2:**
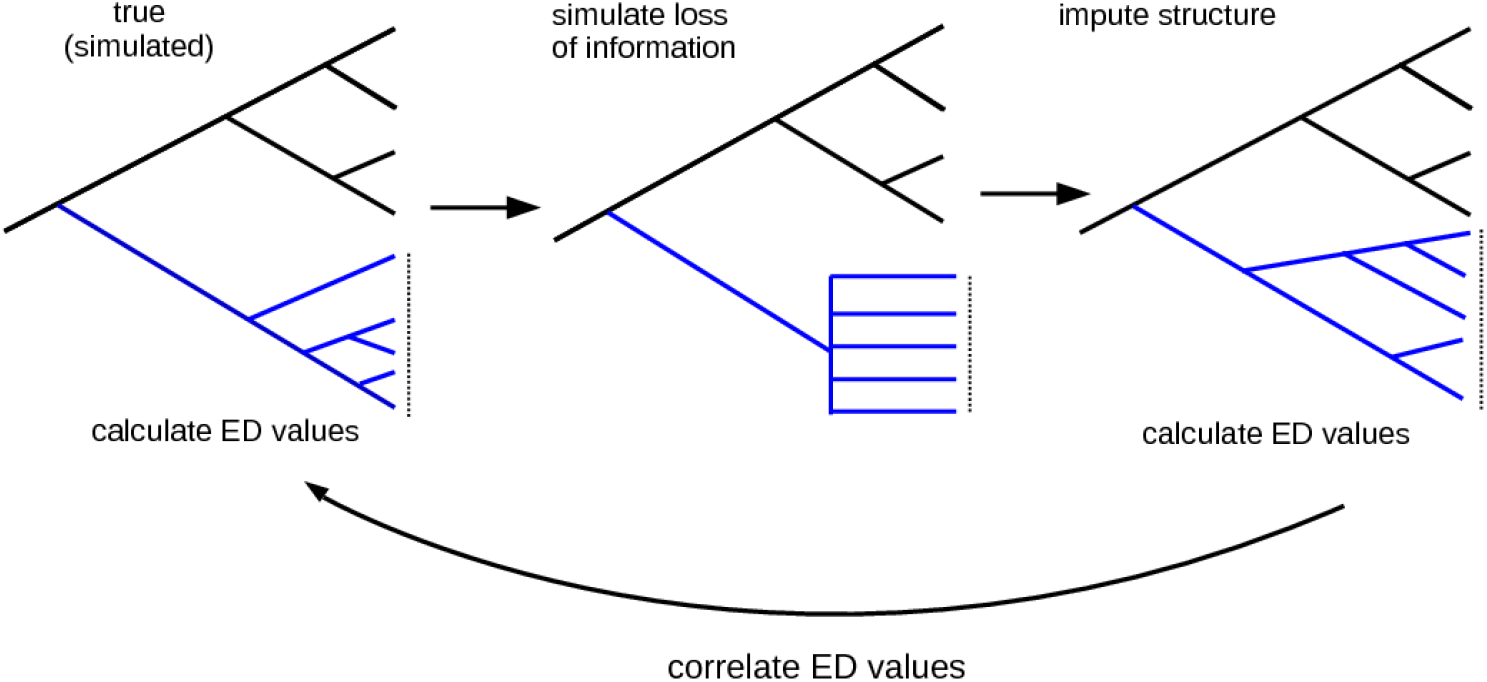
Conceptual overview of the imputation simulations conducted in this study. The simulated tree on the left is the ‘true tree’. We selected a clade to treat as ‘missing’ (highlighted with a dashed line and in blue) by treating it as a polytomy (middle panel), and then imputed the ‘missing’ species to produce the imputed clade in the right panel. To compare true and imputed ED values within the imputed clade, we correlated ED values calculated for the true clade (left) with those for the imputed clade (right).

To assess whether clades, once imputed, had similar ED scores to their true values, we correlated the imputed ED scores with the true ED scores. We also calculated the sum of the absolute change in ranked ED for all species, which is particularly relevant for EDGE-listing as conservation actions are often focused around the top 100, 200, etc., species. Moreover, the correlation of imputed and real scores are bounded by the depth of the imputed clade, and therefore a high correlation could still produce inaccurate imputed scores, and a low correlation could still not be important (*e.g.* they could be anticorrelated but still differ in rank by a max of the size of the subclade). We modeled both of these metrics (the change in ranking and the correlation) as a function of a number of potential explanatory variables. Specifically, we included in our models: the estimated speciation rate of the original phylogeny (using ‘ape::yule’; Paradis, Claude, & Strimmer, 2004), the sum of all phylogenetic branch-lengths in the original phylogeny (Faith’s PD; Faith, 1992), the sum of all phylogenetic branch-lengths in the original focal clade (Faith’s PD; Faith, 1992), the value of γ in the original phylogeny (using ‘phytools::gammatest’; Pybus & Harvey, 2000; Revell, 2012), Colless’ index of the original phylogeny (using ‘apTreeshape::as.treeshape’; Bortolussi, Durand, Blum, & Blum, 2009; Colless, 1982), the kurtosis of species’ ED values in the original phylogeny (using ‘moments::kurtosis’; Komsta & Novomestky, 2015), the skew of species’ ED values in the original phylogeny (using ‘moments::skew’; Komsta & Novomestky, 2015), the total number of species in the original phylogeny, the total number of species within the imputed clade, and the depth (age) of the imputed clade in the phylogeny. Although the expectations of many of these explanatory variables are known for Yule trees, in each simulation they are expected to vary somewhat by chance.

Recently, there has been interest in assigning missing species the mean ED score of the most exclusive clade which contain the species (see Gumbs et al., 2018). To test the efficacy of such methods, we assigned the average ED of the selected clade to each of its’ species and calculated (as above) the mean change in absolute ranking under this scheme. Note that we could not correlate ED scores (as we do above), since such a correlation would require variation in species’ scores and under this approach a single score (the mean ED) is assigned to all imputed species.

## Results

We asked how robust ED scores were for species with known positions on the phylogeny, when other species were missing from the phylogeny. In fact, when there were increasing numbers of missing species, ED scores for the remaining species’ became less accurate (table 1; figure 3). When species were missing from the tree in a phylogenetically-based fashion, ED values were less robust as compared to when species are randomly missing from the tree. However, the effect of missing species is not necessarily severe; even if 20% of species are missing from the tree, the average correlation coefficient between true and estimated ED scores for the remaining species is 0.88 and 0.94 for phylogenetically-biased and random missing species, respectively.

We also considered the impact of imputation on the accuracy of ED scores for imputated species. When clades were imputed on the tree, we found a weak (if any) average positive correlation between the imputed ED and true ED values for species within the imputed clades (overall mean correlation of 0.197 in a statistical model with an r^2^ of 0.5%; figure 4, table 2). We also found no explanatory variables that explained significant variation in this relationship (table 2; see Appendix S1 in Supporting Information). However, we did find evidence that, when imputing larger clades, the variation in the correlation between true and imputed ED scores decreases, although we emphasise the effect is weak (see table 2). When considering rankings rather than raw scores, we found that imputation can introduce sizable error into the estimation of species’ ED values (figure 5 and table 3). This ranking error increased with the size of the imputed clade and phylogeny (table 3), and can affect ranking error within the top 100 and 250 species (see Appendix S2 in Supporting Information). To give an example of the magnitude of the effect, within a phylogeny of 1024 species, the members of an imputed clade of 30 species are, on average, ± 315 rankings from their true rankings. We found similar effects in ranking error when using the average ED value of clade for a missing species (see Appendix S3 in Supporting Information).

**Table 1:** Statistical model of the effect of missing data on the calculation of the remaining species’ ED values. Results of a multiple regression fit to the data shown in figure 3, regressing the correlation coefficient of (remaining) species’ ED scores before and after other species were removed from the phylogeny (*F*_139696,5_ = 40, 350, *r*^2^ = 0.5908, *p* < 0.0001). We emphasize that these are simulated data, and so, as the extremely large sample sizes are likely driving the low standard errors of the model terms, we encourage the reader to focus on the magnitudes of the effects and the overall variance explained by our model (*r*^2^ = 0.5908).The first three rows refer to the overall intercept, effect of the fraction of species removed from the phylogeny, and the overall size of the phylogeny when species were removed in a phylogenetically biased fashion. The last three rows are contrasts, reporting the difference (contrast) of each parameter when species were removed at random from the phylogeny. whether random loss of species has a statistically different effect. The correlation of ED scores appears affected by an interaction between the number of species removed from the tree and whether those species were removed at random or in a phylogenetically-biased fashion. The overall size of the phylogeny has little discernible effect, and its statistical significance is likely driven by the large number of simulations we performed (139, 700).

|  | Estimate | Std. Error | t value | Pr(>\|t\|) |
| --- | --- | --- | --- | --- |
| Reference—Phylogenetically biased |  |  |  |  |
| Intercept | 1.0315 | 0.0013 | 821.39 | <0.0001 |
| Fraction of species removed | -0.4696 | 0.0020 | -233.16 | <0.0001 |
| Number of species overall | $2.500 \times 10^{-6}$ | $2.984 \times 10^{-7}$ | 7.89 | <0.0001 |
| Contrast—Random |  |  |  |  |
| Intercept | 0.0630 | 0.0018 | 35.47 | <0.0001 |
| Fraction of species removed | -0.2774 | 0.0028 | -97.45 | <0.0001 |
| Number of species overall | $5.013 \times 10^{-6}$ | $4.219 \times 10^{-7}$ | -4.38 | <0.0001 |

**Table 2:** Statistical model of the potential drivers of the correlation between imputed and true ED values. Results of a multiple regression fitted to the data shown in figure 4, showing a relatively poor correlation between imputed and true ED scores (*F*_44791,8_ = 29.1, *r*^2^ = 0.005, *p* < 0.0001). Given the extremely low predictive power of this statistical model we are reticent to make strong claims about drivers of the correlation between imputed and observed ED.

|  | Estimate | Std. Error | t value | Pr(>\|t\|) |
| --- | --- | --- | --- | --- |
| Intercept | 0.1974 | 0.0501 | 3.94 | 0.0001 |
| Size of Focal Clade | -0.0036 | 0.0005 | -7.60 | < 0.0001 |
| Size of Phylogeny | 0.0001 | 0.0001 | 0.60 | 0.5497 |
| PD | -0.0001 | 0.0001 | -0.64 | 0.5241 |
| Estimated speciation rate | -0.0199 | 0.0493 | -0.40 | 0.6865 |
| Colless’ Index | -0.0000 | 0.0000 | -0.08 | 0.9380 |
| Skew | 0.0022 | 0.0083 | 0.27 | 0.7885 |
| Kurtosis | -0.0001 | 0.0008 | -0.16 | 0.8736 |
| Depth of Imputed Clade | 0.0006 | 0.0005 | 1.27 | 0.2045 |

**Table 3:** Statistical mode of the effect of clade and phylogeny size on ranking error. Model of the raw data underlying figure 5, regressing the ranking error of imputed species against the number of species in the imputed clade and the entire phylogeny (*F*_47997,2_ = 77890, *r*^2^ = 0.7644, *p* < 0.0001). As can be seen in figure 5, the average ranking error is positively correlated with the size of the clade being imputed and the entire phylogeny. Square-root transformations have been applied to both ranking error and size of phylogeny.

|  | Estimate | Std. Error | t value | Pr(>\|t\|) |
| --- | --- | --- | --- | --- |
| Intercept | -1.6344 | 0.0332 | -49.29 | 0.0001 |
| Size of focal (imputed) clade | 0.0900 | 0.0010 | 91.22 | <0.0001 |
| Size of phylogeny | 0.5179 | 0.0013 | 383.99 | <0.0001 |

**Figure 3:**
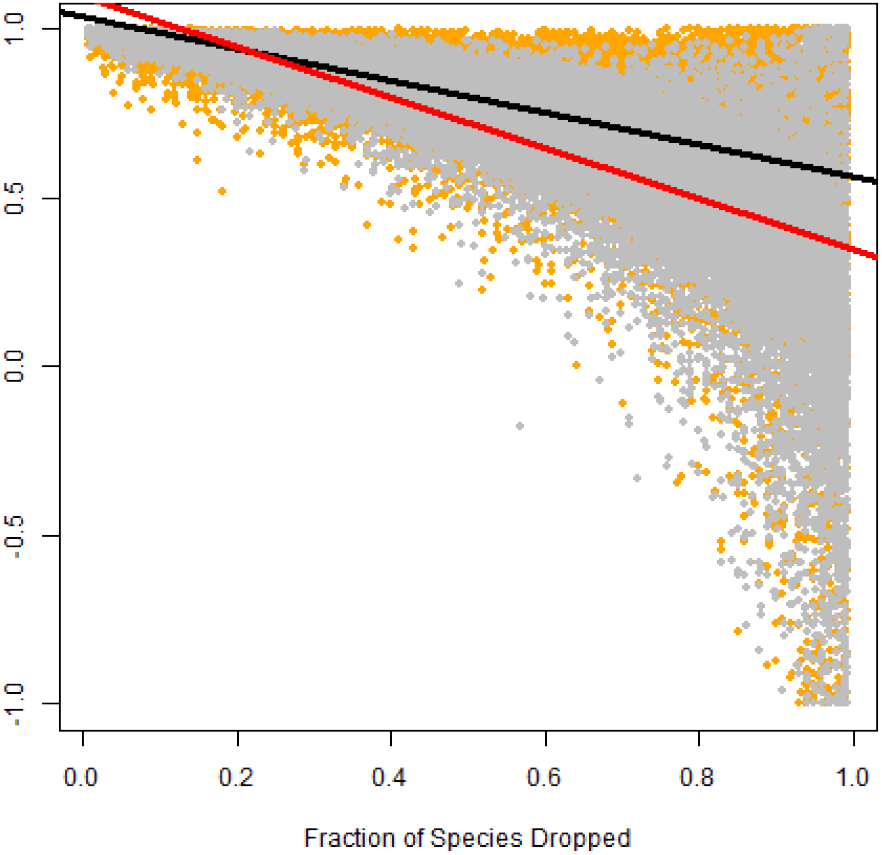
The effect of missing data on the calculation of the remaining species’ ED values. The correlation coefficient of species’ ED values in full (simulated) phylogenies, comparing values before and after the random loss of (other) species from the tree. The color of data points denote whether the species were removed from the phylogeny completely at random (orange) or in a phylogenetically biased fashion (see text; grey). Lines show regressions for random (red) or phylogenetically biased (black) species loss; see table 1 for model coefficients. This plot shows that the accuracy of estimation of ED values is inversely proportional to the number of species missing from the phylogeny, and that phylogenetically-biased species loss has a greater impact on accuracy.

**Figure 4:**
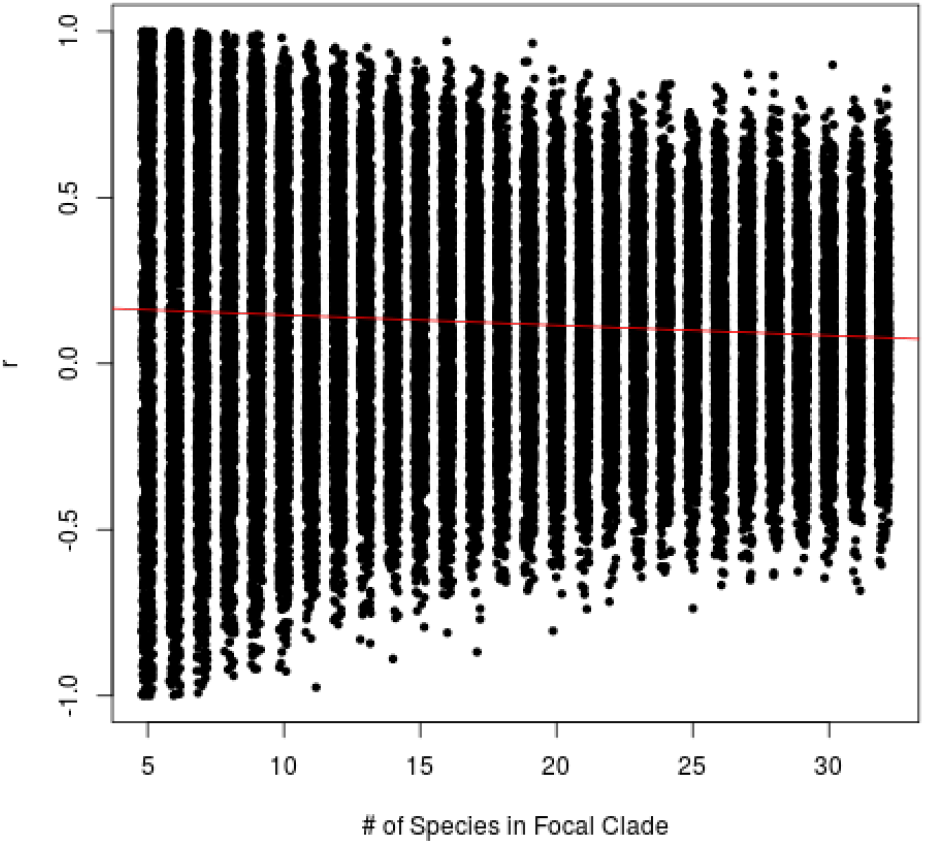
The correlation between species’ imputed and true ED scores plotted as a function of the number of species imputed (focal clade size from all sizes of phylogenies used (n = 128, …, 1024)). Each data point represents the correlation between ED values within the focal clades where imputation has occurred, comparing species’ true ED values with their imputed ED values. This plot, and the statistical analysis of it in table 2, show limited support for an association between true and imputed ED values.

**Figure 5:**
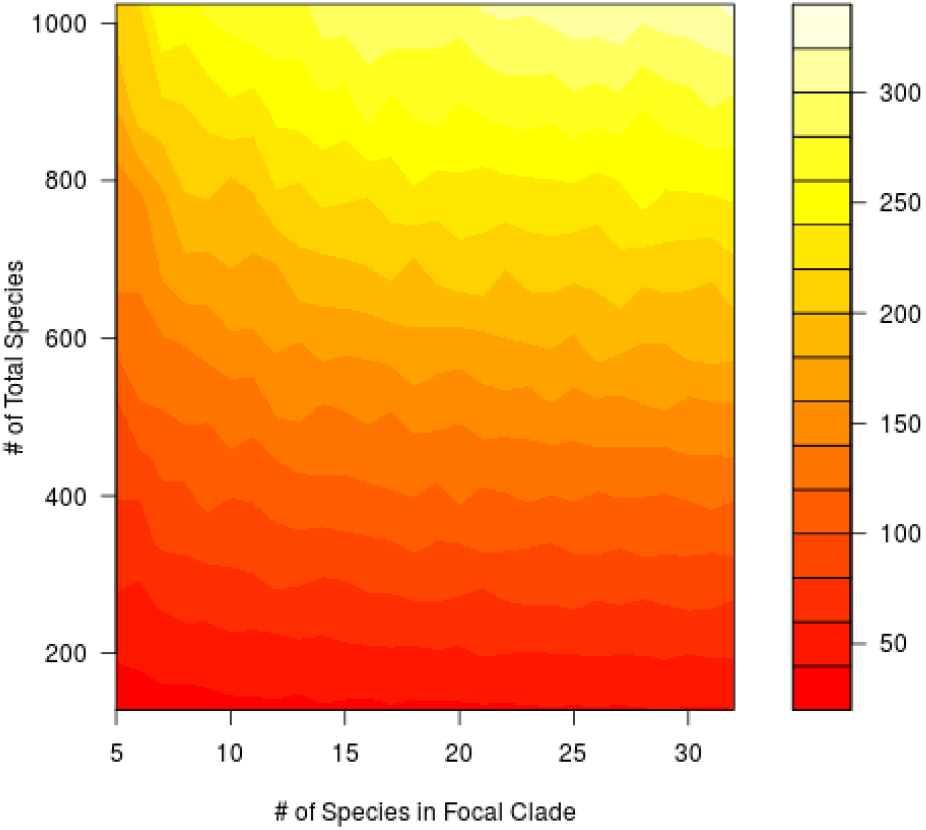
Mean ranking error of imputed species. An interpolated heat-map of the mean ranking error of imputed species as a function of the total number of species in the phylogeny (vertical axis) and number of species in the focal (imputed) clade (horizontal axis). Table 3 gives statistical support for the trend of increased error in larger phylogenies and imputed clades.

## Discussion

Phylogenies are playing an increasing role in conservation prioritization, decision-making, and policy (Veron, Davies, Cadotte, Clergeau, & Pavoine, 2017; Vézquez & Gittleman, 1998). A major obstacle to a more widespread adoption of phylogenetic prioritization methods such as EDGE is phylogenetic uncertainty (Collen, 2015). There is a tension between a purported need to make decisions to preserve biodiversity— including evolutionary history—now, and the reality that we rarely have complete information about the phylogenetic placement of many species of conservation concern (N. J. Isaac & Pearse, in press). The intention of our study is to provide concrete information about the impact of one source of phylogenetic uncertainty - missing species - on conservation prioritization. To address this uncertainty, we addressed two key issues: (1) the extent to which species that are missing from the tree of life impact the ED scores of species for which we do have data, and (2) the extent to which phylogenetic imputation can accurately estimate ED scores for taxa with no phylogenetic data. First, we found that missing species had a surprisingly small impact on the ED scores of other species, particularly if species are missing at random from the tree of life. Second, we found that phylogenetic imputation generally fails to accurately reconstruct species’ ED scores and rankings.

Our results are derived solely from simulations under a simple model of diversification—the Yule model. We acknowledge that lineages evolve in more complex ways. However, such complexities are unlikely to make imputation easier, and they would have to be modelled and quantified in empirical studies. We suggest that focusing on the simplest model of diversification makes our results more generalizable. Further, we focus here solely on the results from a single imputation in each simulation, despite, empirically, biologists reporting average ED scores calculated across pseudo-posterior distributions of many imputed phylogenies (Kuhn et al., 2011). Thus our results show that the variation within these pseudo-posterior distributions is likely very large. It is well-known that such imputation methods are not biased (indeed, this was originally shown by Kuhn et al., 2011): here we emphasize that the uncertainty they introduce is sufficiently large such that they may be less informative than previously has been thought.

### ED scores are relatively robust to missing species

Missing species and poor phylogenetic resolution have been identified as causes of uncertainty when calculating ED (Isaac et al., 2007), but we were unable to find a quantitative assessment of how missing species might affect ED values of species for which data is available. Empirically in corals and gymnosperms, incomplete phylogenies produced similar results as later, more complete trees (Curnick et al., 2015; Forest et al., 2018). Our results support this finding. Indeed, our analysis suggests that, on average (and we emphasize that there is a good amount of variation about that average; see figure 3), a phylogeny missing 20% of species at random will still have ED scores for the remaining species that are strongly correlated (mean rho = 0.94) with the true ED scores.

We did find that missing species are more problematic when those species are non-randomly distributed across the phylogeny. Our simulations do not examine extreme phylogenetic patterning, such as if an entire clade were missing. This is notable because clades that are geographically restricted to difficult-to-reach regions are both difficult to sequence and not uncommon (as is seen with 27 coral species in the Indian Ocean; Arrigoni, Stefani, Pichon, Galli, & Benzoni, 2012). We also do not attempt to comprehensively simulate all of the different ways in which species could be missing from a phylogeny. We simply demonstrate that, compared to a scenario in which species are missing at random, phylogenetically patterned missing species can have a greater effect on the ED scores of species for which we have data.

### Imputation does not reconstruct the ED values of missing species with great precision

Our results show that neither imputation (figures 4 and 5), nor clade-averages of ED (see Appendix S3 in Supporting Information), accurately recover the true ED values or the true ED rank of missing species. Thus we argue that, even though imputation allows missing species to be incorporated into EDGE lists, their associated EDGE scores may not accurately reflect their true scores. We acknowledge these are averages and may change depending on particular phylogeny, but we can find no statistically significant predictors of that variation.

While we did not assess clades with fewer than five species (we do not consider correlations or averages to be reliable with so few data-points), we cannot think why smaller clades would necessarily be more reliable (and this would require a large deviation from the trend in figure 4). Indeed, in the smallest possible clade (two species), imputation is essentially sampling a terminal branch length from an exponential distribution (Kuhn et al., 2011); such a process should still lead to a great degree of uncertainty.

It is, perhaps, unsurprising that imputed ED values do not correlate with their true values (see figure 4), but we were surprised at the degree of ranking error. Indeed, larger phylogenies showed *greater* ranking error; we naïvely would have expected the opposite. We would expect the the upper bound on the age of the imputed clade, which should have expected be relatively younger in larger phylogenies, would partially controlled the range of the ranks for the imputed species. ED is known to be driven mostly by terminal branch length (Isaac et al., 2007; D. W. Redding et al., 2008; Steel et al., 2007); our results therefore emphasize this.

Imputation is not the only way to incorporate missing species into EDGE-like frameworks (see Collen et al., 2011; Gumbs et al., 2018), but it is likely the most common. 3, 330 of the birds (^~^30%; Jetz et al., 2014), 250 of the mammals (^~^5.6%; Collen et al., 2011), and 610 of the sharks (^~^49%; Stein et al., 2018) in recent EDGE lists were imputed. It is well-known that phylogenetic imputation can cause biases in other statistical methods, such as the estimation of evolutionary phylogenetic signal (Rabosky, 2015). We emphasize that we are not suggesting that imputation *biases* ED scores: we are, instead, suggesting that it is less precise than has previously been acknowledged.

### Guidelines for the use of imputation

The impact of imputation on EDGE scores is almost certainly less than its impact on ED scores, because EDGE scores are a product of both ED and IUCN status (‘GE’). However, the goal of EDGE-like measures is to incorporate phylogeny, and if imputed EDGE scores are driven by their GE component because of uncertainty introduced by imputation, this essentially creates another metric of IUCN status.

Our results further suggest that incomplete phylogenies can be used to estimate ED scores with remarkably high degrees of accuracy. Instead of using imputation to account for the relatively minor impact of missing species, we suggest that conservation biologists should, without accounting for phylogenetic uncertainty, focus on the species for which they have data. While we have not explored this uncertainty here, evolutionary biologists commonly work with distributions of trees generated from genetic data (reviewed in Bollback, 2005; Huelsenbeck, Ronquist, Nielsen, & Bollback, 2001), since the precise topology and dating of a phylogeny is almost always uncertain. This uncertainty has, indeed, already been shown to affect EDGE scores and rankings (Pearse et al., 2015). If biologists are concerned about the impact of missing species on known species’ ED(GE) scores we see no harm in being precautionary and using imputation. It is important, however, to focus on known sources of potential error, and so we would encourage biologists to incorporate uncertainty in species with phylogenetic data as a priority.

Our results suggest that prioritizing species whose phylogenetic structure has been imputed should be done with extreme care, if at all. In the case that an species is imputed to be below a threshold set for conservation (most EDGE studies focus on the ‘top 100’ species or something similar), then the path forward is clear: that species should not have conservation funds allocated to it at this time. The case where a species, on average, passes a threshold is more complex, but the theory underlying imputation can give some guidance. Imputed distributions of trees essentially represent Bayesian posterior distributions (Kuhn et al., 2011), and so the 95% posterior densities of these distributions’ ED values represent a range within which we can be 95% certain the true ED scores lie (if the model assumptions are met). Thus we suggest that conservation action should only be initiated for a species if there is a 95% (or 80%, or whatever confidence is deemed appropriate) probability that it is above that threshold. For example, a species whose ranking is estimated to have a 20% probability of being between the 1^st^ and 100^th^ highest-ranked species could not, with confidence, be called a top-100 species. Our results suggest that, on average, very few imputed species will meet such a criterion. Regardless, the calculations of such probabilities is trivial with the data users of imputation have in hand already.

Ultimately, we are currently fighting a losing battle to preserve the tree of life. Our results are good news: they suggest that we can start right away using the (incomplete) phylogenies we already have. The effect of missing species is negligible enough that we often do not need time-consuming imputation, and imputation rarely gives us sufficiently precise estimates of species’ ED scores anyway. We suggest that, given we do not have the resources to save everything, we should consider focusing our efforts on those species whose ED scores we can know with greater certainty: those for which we have data.

## Acknowledgments

We thank the Department of Biology and Ecology Center of Utah State University for their support. We are grateful to E. Simpson, M. Sneddon, J. Stachewicz and J. Rosindell for ongoing discussion and-or constructive feedback on this manuscript.

## Data Accessibility Statement

All simulation and analysis code, along with underlying data, generated for this study are in the supplementary materials and online at: url https://github.com/bweedop/edgeSims

## References

Arrigoni, R., Stefani, F., Pichon, M., Galli, P., & Benzoni, F. (2012). Molecular phylogeny of the robust clade (faviidae, mussidae, merulinidae, and pectiniidae): An indian ocean perspective. Molecular Phylogenetics and Evolution, 65(1), 183–193.

Barnosky, A. D., Matzke, N., Tomiya, S., Wogan, G. O., Swartz, B., Quental, T. B., … Maguire, K. C. (2011). Has the earth’s sixth mass extinction already arrived? Nature, 471(7336), 51–57.

Bollback, J. (2005). Posterior mapping and posterior predictive distributions. In R. Neilsen (Ed.), Statistical methods in molecular evolution (Chap. 16, pp. 439–462). Springer New York.

Bortolussi, N., Durand, E., Blum, M., & Blum, M. M. (2009). Package ‘aptreeshape’.

Bottrill, M. C., Joseph, L. N., Carwardine, J., Bode, M., Cook, C., Game, E. T., … McDonald-Madden, E. (2008). Is conservation triage just smart decision making? Trends in Ecology & Evolution, 23(12), 649–654.

Brooks, T. M., Mittermeier, R. A., Mittermeier, C. G., Da Fonseca, G. A., Rylands, A. B., Konstant, W. R., … Magin, G. (2002). Habitat loss and extinction in the hotspots of biodiversity. Conservation biology, 16(4), 909–923.

Butchart, S. H., & Bird, J. P. (2010). Data deficient birds on the iucn red list: What don’t we know and why does it matter? Biological Conservation, 143(1), 239–247.

Ceballos, G., Ehrlich, P. R., Barnosky, A. D., García, A., Pringle, R. M., & Palmer, T. M. (2015). Accelerated modern human-induced species losses: Entering the sixth mass extinction. Science advances, 1(5), e1400253.

Collen, B. (2015). Conservation prioritization in the context of uncertainty. Animal Conservation, 18(4), 315–317.

Collen, B., Turvey, S. T., Waterman, C., Meredith, H. M., Kuhn, T. S., Baillie, J. E., & Isaac, N. J. (2011). Investing in evolutionary history: Implementing a phylogenetic approach for mammal conservation. Philosophical Transactions of the Royal Society of London B: Biological Sciences, 366(1578), 2611–2622.

Colless, D. H. (1982). Review of phylogenetics: The theory and practice of phylogenetic systematics. Systematic Zoology, 31(1), 100–104.

Curnick, D., Head, C., Huang, D., Crabbe, M. J. C., Gollock, M., Hoeksema, B., … Obura, D. (2015). Setting evolutionary-based conservation priorities for a phylogenetically data-poor taxonomic group (scleractinia). Animal Conservation, 18(4), 303–312.

Dulvy, N. K., Fowler, S. L., Musick, J. A., Cavanagh, R. D., Kyne, P. M., Harrison, L. R., … Francis, M. P. (2014). Extinction risk and conservation of the world’s sharks and rays. Elife, 3.

Faith, D. P. (1992). Conservation evaluation and phylogenetic diversity. Biological conservation, 61(1), 1–10.

Felsenstein, J. (2005). Using the quantitative genetic threshold model for inferences between and within species. Philosophical Transactions of the Royal Society of London B: Biological Sciences, 360(1459), 1427–1434.

Forest, F., Moat, J., Baloch, E., Brummitt, N. A., Bachman, S. P., Ickert-Bond, S., … Mathews, S. (2018). Gymnosperms on the edge. Scientific reports, 8(1), 6053.

Good, T. C., Zjhra, M. L., & Kremen, C. (2006). Addressing data deficiency in classifying extinction risk: A case study of a radiation of bignoniaceae from madagascar. Conservation Biology, 20(4), 1099–1110.

Gumbs, R., Gray, C. L., Wearn, O. R., & Owen, N. R. (2018). Tetrapods on the edge: Overcoming data limitations to identify phylogenetic conservation priorities. PloS One, 13(4), e0194680.

Huelsenbeck, J. P., Ronquist, F., Nielsen, R., & Bollback, J. P. (2001). Bayesian inference of phylogeny and its impact on evolutionary biology. Science, 294(5550), 2310–2314.

Isaac, N. J., Redding, D. W., Meredith, H. M., & Safi, K. (2012). Phylogenetically-informed priorities for amphibian conservation. PLoS one, 7(8), e43912.

Isaac, N. J., Turvey, S. T., Collen, B., Waterman, C., & Baillie, J. E. (2007). Mammals on the edge: Conservation priorities based on threat and phylogeny. PloS one, 2(3), e296.

IUCN. (2001). Iucn red list categories and criteria. IUCN.

IUCN. (2008). Guidelines for using the iucn red list categories and criteria, version 7.0. Prepared by the Standards and Petitions Working Group of the IUCN SSC Biodiversity Assessments Sub-Committee in August 2008.

Jensen, E. L., Mooers, A. Ø., Caccone, A., & Russello, M. A. (2016). I-hedge: Determining the optimum complementary sets of taxa for conservation using evolutionary isolation. PeerJ, 4, e2350.

Jetz, W., Thomas, G. H., Joy, J. B., Redding, D. W., Hartmann, K., & Mooers, A. O. (2014). Global distribution and conservation of evolutionary distinctness in birds. Current Biology, 24(9), 919–930.

Komsta, L., & Novomestky, F. (2015). Moments, cumulants, skewness, kurtosis and related tests. R package version 0.14.

Kuhn, T. S., Mooers, A. Ø., & Thomas, G. H. (2011). A simple polytomy resolver for dated phylogenies. Methods in Ecology and Evolution, 2(5), 427–436.

Lips, K. R., Brem, F., Brenes, R., Reeve, J. D., Alford, R. A., Voyles, J., … Collins, J. P. (2006). Emerging infectious disease and the loss of biodiversity in a neotropical amphibian community. Proceedings of the national academy of sciences of the United States of America, 103(9), 3165–3170.

McBride, M. F., Wilson, K. A., Bode, M., & Possingham, H. P. (2007). Incorporating the effects of socioeconomic uncertainty into priority setting for conservation investment. Conservation Biology, 21(6), 1463–1474.

Molnar, J. L., Gamboa, R. L., Revenga, C., & Spalding, M. D. (2008). Assessing the global threat of invasive species to marine biodiversity. Frontiers in Ecology and the Environment, 6(9), 485–492.

Morais, A. R., Siqueira, M. N., Lemes, P., Maciel, N. M., De Marco Jr, P., & Brito, D. (2013). Unraveling the conservation status of data deficient species. Biological conservation, 166, 98–102.

Myers, N., Mittermeier, R. A., Mittermeier, C. G., Da Fonseca, G. A., & Kent, J. (2000). Biodiversity hotspots for conservation priorities. Nature, 403(6772), 853–858.

Naidoo, R., Balmford, A., Ferraro, P. J., Polasky, S., Ricketts, T. H., & Rouget, M. (2006). Integrating economic costs into conservation planning. Trends in ecology & evolution, 21(12), 681–687.

Orme, D., Freckleton, R., Thomas, G., Petzoldt, T., Fritz, S., Isaac, N. J., & Pearse, W. D. (2013). Caper: Comparative analyses of phylogenetics and evolution in r. R package version 0.5.2. Retrieved from https://CRAN.R-project.org/package=caper

Paradis, E., Claude, J., & Strimmer, K. (2004). APE: Analyses of phylogenetics and evolution in R language. Bioinformatics, 20, 289–290.

Pearse, W. D., Chase, M. W., Crawley, M. J., Dolphin, K., Fay, M. F., Joseph, J. A., … Roy, D. B. (2015). Beyond the edge with edam: Prioritising british plant species according to evolutionary distinctiveness, and accuracy and magnitude of decline. PloS one, 10(5), e0126524.

Pennell, M. W., Eastman, J. M., Slater, G. J., Brown, J. W., Uyeda, J. C., FitzJohn, R. G., … Harmon, L. J. (2014). Geiger v2. 0: An expanded suite of methods for fitting macroevolutionary models to phylogenetic trees. Bioinformatics, 30(15), 2216–2218.

Pounds, J. A., Bustamante, M. R., Coloma, L. A., Consuegra, J. A., Fogden, M. P., Foster, P. N., … Puschendorf, R. (2006). Widespread amphibian extinctions from epidemic disease driven by global warming. Nature, 439(7073), 161–167.

Pressey, R., Humphries, C., Margules, C. R., Vane-Wright, R., & Williams, P. (1993). Beyond opportunism: Key principles for systematic reserve selection. Trends in ecology & evolution, 8(4), 124–128.

Pybus, O. G., & Harvey, P. H. (2000). Testing macro-evolutionary models using incomplete molecular phylogenies. Proceedings of the Royal Society of London B: Biological Sciences, 267(1459), 2267–2272.

R Core Team. (2017). R: A language and environment for statistical computing. R Foundation for Statistical Computing. Vienna, Austria. Retrieved from https://www.R-project.org/

Rabosky, D. L. (2015). No substitute for real data: A cautionary note on the use of phylogenies from birth-death polytomy resolvers for downstream comparative analyses. Evolution, 69(12), 3207–3216.

Redding, D. (2003). Incorporating genetic distinctness and reserve occupancy into a conservation priorisation approach. Master’s thesis, University of East Anglia, Norwich OpenURL.

Redding, D. W., Hartmann, K., Mimoto, A., Bokal, D., DeVos, M., & Mooers, A. Ø. (2008). Evolutionarily distinctive species often capture more phylogenetic diversity than expected. Journal of theoretical biology, 251(4), 606–615.

Revell, L. J. (2012). Phytools: An r package for phylogenetic comparative biology (and other things). Methods in Ecology and Evolution, 3, 217–223.

Steel, M., Mimoto, A., & Mooers, A. Ø. (2007). Hedging our bets: The expected contribution of species to future phylogenetic diversity. Evolutionary bioinformatics online, 3, 237.

Stein, R. W., Mull, C. G., Kuhn, T. S., Aschliman, N. C., Davidson, L. N., Joy, J. B., … Mooers, A. O. (2018). Global priorities for conserving the evolutionary history of sharks, rays and chimaeras. Nature Ecology & Evolution, 1.

Thomas, C. D., Cameron, A., Green, R. E., Bakkenes, M., Beaumont, L. J., Collingham, Y. C., … Hannah, L. (2004). Extinction risk from climate change. Nature, 427(6970), 145–148.

Thomas, G. H., Hartmann, K., Jetz, W., Joy, J. B., Mimoto, A., & Mooers, A. O. (2013). Pastis: An r package to facilitate phylogenetic assembly with soft taxonomic inferences. Methods in Ecology and Evolution, 4(11), 1011–1017.

Tonini, J. F. R., Beard, K. H., Ferreira, R. B., Jetz, W., & Pyron, R. A. (2016). Fully-sampled phylogenies of squamates reveal evolutionary patterns in threat status. Biological Conservation, 204, 23–31.

Veron, S., Davies, T. J., Cadotte, M. W., Clergeau, P., & Pavoine, S. (2017). Predicting loss of evolutionary history: Where are we? Biological Reviews, 92(1), 271–291.

Vézquez, D. P., & Gittleman, J. L. (1998). Biodiversity conservation: Does phylogeny matter? Current Biology, 8(11), R379–R381.

Wake, D. B., & Vredenburg, V. T. (2008). Are we in the midst of the sixth mass extinction? a view from the world of amphibians. Proceedings of the National Academy of Sciences, 105(Supplement 1), 11466–11473.

Weitzman, M. L. (1998). The noah’s ark problem. Econometrica, 1279–1298.

Wilson, K. A., Underwood, E. C., Morrison, S. A., Klausmeyer, K. R., Murdoch, W. W., Reyers, B., … McBride, M. F. (2007). Conserving biodiversity efficiently: What to do, where, and when. PLOS biology, 5(9), e223.

